# Integrated Spatial Metabolomics and Proteomics from the Same Tissue Section Using a Conductive ITO-PET Slide

**DOI:** 10.64898/2026.09.07.749981

**Authors:** Haizhu Wang, Shengmei Liu, Lixian Lin, Yongsheng Cheng, Yuchuang Zheng, Yan Li, Peng Xue

**Affiliations:** Key Laboratory of Epigenetic Regulation and Intervention, Institute of Biophysics, Chinese Academy of Sciences, 15 Datun Road, Beijing 100101, China; Department of Medical-Engineering Integration and Technology, Guangzhou Laboratory, 9 Xingdao Ring North Road, Guangzhou International Bio Island, Haizhu District, Guangzhou 510005, China; The Air Force Hospital of the Southern Theater Command of the Chinese People’s Liberation Army, 801 Dongfeng East Road, Yuexiu District, Guangzhou 510062, Guangdong, China

**Keywords:** Spatial multi-omics, MALDI mass spectrometry imaging, Laser capture microdissection

## Abstract

Integrating spatial metabolomics and spatial proteomics on the same tissue section remains challenging because matrix-assisted laser desorption/ionization mass spectrometry imaging (MALDI-MSI) and laser capture microdissection (LCM)-based proteomics impose different requirements on sample slides. Here, we developed and systematically evaluated a conductive indium tin oxide-coated polyethylene terephthalate (ITO-PET) slide that enables sequential MALDI-MSI and LCM-liquid chromatography-mass spectrometry (LCM-LC-MS) analysis of the same tissue section. Using mouse brain tissue as a model, ITO-PET provided MALDI-MSI performance closely comparable to conventional ITO-glass, including spectral concordance (Pearson correlation, R = 0.90), ion detection coverage, metabolite annotation, signal intensity distribution, and preservation of spatial molecular patterns. Following MALDI-MSI, the ITO-PET slide enabled cutting-mode LCM and yielded proteomic signal intensities and numbers of identified protein groups comparable to those obtained with conventional PEN-glass slides. Across different tissue sampling areas, proteomic signal intensity distributions, precursor ion counts, and protein group identifications remained broadly comparable before and after MALDI-MSI, with substantial overlap in identified protein groups. Similar patterns were observed in mouse kidney, lung, spleen, and liver tissues, further supporting the applicability of the workflow across different tissue types. By combining the electrical conductivity required for MALDI-MSI with the mechanical properties required for LCM cutting, the ITO-PET slide addresses a major material incompatibility between the two analytical modalities and enables sequential spatial metabolomic and proteomic analysis from the same tissue section. This workflow provides a practical analytical platform for obtaining complementary molecular information from spatially limited biological specimens.

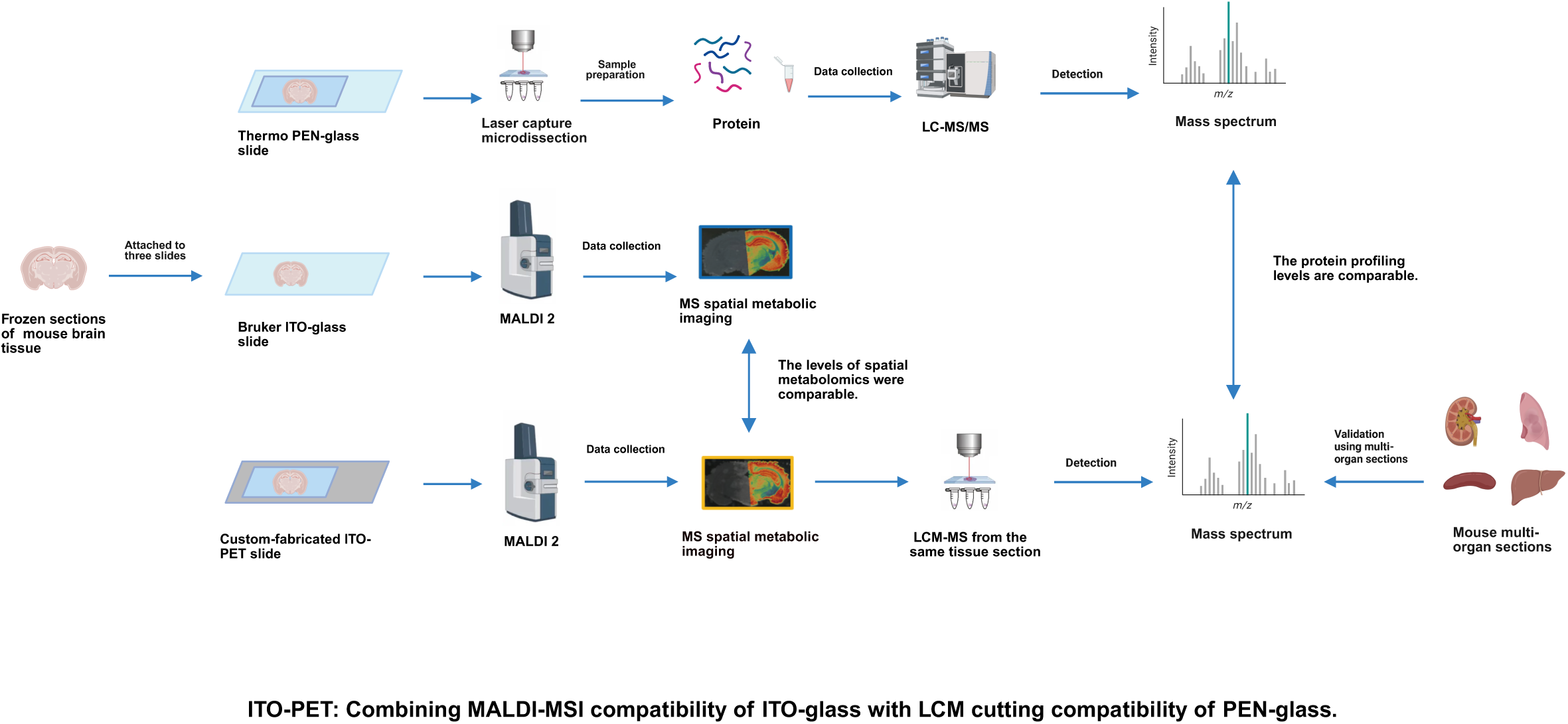

## Introduction

Spatial multi-omics integrates multiple molecular layers, including the genome, transcriptome, proteome, and metabolome, while preserving their native spatial context within tissues.^1–3^ By overcoming the loss of spatial information inherent to conventional bulk omics, spatial multi-omics enables cellular and subcellular localization of molecules and characterization of their spatial distributions and interactions.^4, 5^ These capabilities provide new opportunities to investigate cellular microenvironments,^6, 7^ tissue structure–function relationships,^8^ and the spatiotemporal molecular changes associated with disease.^9^

Among these molecular layers, the metabolome provides a direct readout of cellular biochemical activity and reflects the downstream effects of genomic, epigenetic, transcriptional, and environmental regulation.^10^ Spatial metabolomics therefore provides functional information complementary to genomic and transcriptomic maps.^11^ Matrix-assisted laser desorption/ionization mass spectrometry imaging (MALDI-MSI) is a major platform for spatial metabolomics,^12^ enabling label-free, in situ mapping of metabolites, lipids, and other small molecules on tissue sections at micrometer-scale spatial resolution.^13, 14^ However, a comprehensive understanding of spatial metabolic phenotypes also requires information on the proteins and enzymes that regulate the underlying metabolic pathways. Spatial proteomics provides this complementary molecular information, and laser capture microdissection (LCM) coupled with liquid chromatography–tandem mass spectrometry (LC-MS/MS) enables spatially defined cells or tissue regions to be isolated for downstream proteomic characterization.^15–21^ Recent advances, exemplified by Deep Visual Proteomics, which integrates image analysis, automated LCM, and ultrasensitive mass spectrometry, further demonstrate the potential of spatial proteomics for characterizing molecular heterogeneity within tissues.^22–24^

Despite advances in spatial metabolomics and spatial proteomics, integrating MALDI-MSI and LCM-based proteomics on the same tissue section remains challenging because the two techniques require different slide properties. PEN membrane-coated glass slides (PEN-glass) are compatible with cutting-mode LCM but lack the electrical conductivity required for optimal MALDI-MSI, whereas ITO-coated glass slides (ITO-glass) provide sufficient conductivity for MALDI-MSI but cannot be readily cut during laser microdissection.^25, 26^ Several approaches have been explored to address this compatibility challenge. In 2019,^27^ Dewez et al. combined MALDI-MSI and LCM-based molecular analysis on the same tissue section using PEN membrane slides. However, although PEN-glass slides facilitate cutting-mode microdissection, their lack of electrical conductivity can limit MALDI-MSI performance. In 2021,^25^ Mezger et al. demonstrated mass spectrometry spatial-omics on a single conductive slide using an ITO-coated glass slide. Although conventional ITO-glass provides the conductivity required for MALDI-MSI, it cannot be cut during laser microdissection; consequently, tissue collection requires laser ablation, which can adversely affect downstream proteomic recovery.^26^

More recently, Veličković et al.^28^ introduced metal-assisted strategies to improve the MALDI-MSI compatibility of electrically insulating PEN slides, using either conductive copper-tape backing or a thin gold coating for sequential MALDI-MSI and LCM-based proteomic analysis of the same tissue section. Although these approaches advanced single-section spatial multi-omics, they required additional conductive modifications and showed platform-dependent performance: copper-tape-backed PEN performed well on timsTOF but provided reduced metabolite coverage on FTICR, whereas gold coating required an additional sputter-coating step and specialized instrumentation and was associated with greater tissue damage. An alternative strategy is to perform spatial metabolomic and proteomic analyses on consecutive tissue sections mounted on different slide types;^29, 30^ however, this approach loses direct spatial correspondence between molecular layers and may be affected by molecular heterogeneity between adjacent sections.^28, 31^ Together, these limitations underscore the need for a simpler and more broadly applicable approach to integrating MALDI-MSI and LCM-based proteomics.

To address this need, we developed a custom-fabricated ITO-PET slide that combines the electrical conductivity required for MALDI-MSI with the laser-cutting compatibility required for LCM. This design enables sequential MALDI-MSI and cutting-mode LCM from the same tissue section without additional conductive backing or post-section metal coating. Using mouse brain tissue as a model, we systematically benchmarked ITO-PET against conventional ITO-glass and PEN-glass slides for MALDI-MSI, LCM-based tissue collection, and downstream proteomic analysis, with further evaluation across different tissue sampling areas and multiple organ types. The resulting workflow enables complementary spatial metabolomic and proteomic information to be obtained from the same tissue section while maintaining direct spatial correspondence between molecular layers, providing a practical approach for integrated spatial multi-omics of spatially heterogeneous or limited biological specimens.

## 2 Experimental Section

### 2.1 Fabrication and Characterization of ITO-PET Slides

A commercial 5-μm-thick polyethylene terephthalate (PET) film (DuPont, USA) was used as the base material for fabrication of the ITO-PET slide. The PET film was subsequently coated with an indium tin oxide (ITO) layer by a commercial coating service provider (Gengte, China) to produce the conductive ITO-PET film. According to the manufacturer’s specifications, the resulting ITO coating had a sheet resistance of <20 Ω sq^¹^, an average visible-light transmittance of >79%, and a nominal thickness of approximately 100 nm. The surface roughness was characterized using atomic force microscopy (AFM, Bruker Dimension Icon) over a scan area of 10 × 10 µm. Employing the electrical and optical parameters provided by the supplier as quality benchmarks, the film was mounted on a standard-sized metal frame, thoroughly cleaned sequentially with deionized water, methanol, and acetonitrile, and subsequently stored in a dust-free environment for further use.

### 2.2 Preparation of Mouse Brain Tissue Sections on Different Slides

Fresh brain tissue samples were obtained from 10-week-old male BALB/c mice. Animals were housed in a specific pathogen-free facility under a 12-h light/dark cycle at 22 ± 2 °C and 50 ± 10 % humidity, with free access to food and water. All animal procedures were approved by the Institutional Animal Care and Use Committee of Guangzhou Laboratory (approval no. GZLAB-AUCP-202509-A06). Euthanasia was performed by cervical dislocation, and death was confirmed by subsequent decapitation.^32^ Following rapid dissection, the brain tissue was immediately flash frozen in liquid nitrogen without embedding to preserve native molecular integrity. Coronal sections of 10 μm thickness were cut using a cryostat (Leica, Germany) maintained at approximately –18 °C (chamber and knife temperature).^33, 34^ The sections were carefully thaw mounted onto three distinct slides: a commercial ITO coated glass slide (Bruker, Germany), the custom-fabricated ITO PET slide, and a standard PEN (polyethylene naphthalate) membrane slide (Thermo Fisher Scientific, USA). Sections were either analyzed immediately or stored at –80 °C for up to two weeks before further analysis, according to Bruker’s technical support.

### 2.3 MALDI Mass Spectrometry Imaging

Prior to MALDI MSI analysis, all slides were vacuum dried in a desiccator for 20 minutes. A uniform layer of 2,5 dihydroxybenzoic acid (DHB) matrix was deposited onto the tissue sections using an HTX TM Sprayer automated sprayer system (HTX, USA).^35^ The matrix solution consisted of 15 mg/mL DHB in 90:10 (v/v) acetonitrile (ACN)/water containing 0.1% trifluoroacetic acid (TFA), and was applied onto the tissue sections using a syringe pump. Spraying parameters were as follows: nozzle temperature 60 °C; matrix flow rate 0.125 mL/min; nitrogen pressure 10 psi; nozzle velocity 1200 mm/min; nozzle height 40 mm; spray spacing 2 mm; 10 spray(cycles) passes with a 5 s drying time between passes.

MALDI mass spectrometry imaging was performed on a MALDI 2 timsTOF fleX instrument (Bruker, Germany) equipped with microgrid technology.^36, 37^ Before each imaging run, mass calibration was performed using a tuning mix solution (Agilent, USA). The SmartBeam laser was set to “Custom” mode with “Single” enabled, delivering 300 laser shots per pixel at a repetition rate of 10 kHz within a pixel area of 30 × 30 μm². Step sizes were set to 30 μm in both the x and y directions. Data were acquired in positive ion mode over a mass range of m/z 100–1300. The collision cell energy offset was set to 8 eV, the collision radio-frequency (RF) voltage was set to 1200 Vpp, the quadrupole ion energy was set to 5 eV, the low mass cut off was set to m/z 100, and the pre TOF transfer time was 60 μs. The second laser of the MALDI 2 source was activated, while trapped ion mobility separation (TIMS) was disabled (TIMS off) for all imaging experiments.

### 2.4 MSI Data Processing and Analysis

The acquired MSI data from all three substrates were systematically processed using Bruker SCiLS Lab software (version 2023b)Metaspace Online Platform and MetaboScape 2025b. For visualization of the average mass spectra, intensity thresholds of >100 were applied to the ITO-glass and ITO-PET datasets, whereas a lower threshold of >10 was applied to the PEN-glass dataset because of its substantially lower overall signal intensity. All detected m/z features were exported for subsequent comparison without applying an additional intensity threshold. Venn diagrams were used to visualize the unique and shared m/z features among ITO-glass, ITO-PET, and PEN-glass based on the exported feature lists.

For pairwise comparison of ion signal intensities, signals with intensities >1 were retained and log-transformed prior to Pearson correlation analysis and linear regression. For pairwise intensity-ratio visualization, shared m/z features with signal intensities >50 were retained, and intensity ratios were calculated between the corresponding slide pairs.

Spatial organization of the MALDI-MSI datasets was evaluated using Leiden clustering of pixel-wise spectral profiles implemented in Scanpy, with the clustering resolution set to 0.5. Uniform manifold approximation and projection (UMAP) was performed using Scanpy for dimensionality reduction and visualization of the high-dimensional spectral data, with the number of nearest neighbors (n_neighbors) set to 15, the minimum distance (min_dist) set to 0.1, and Euclidean distance used as the metric. The same clustering and dimensionality-reduction parameters were applied across the datasets. Pairwise UMAP comparison between ITO-glass and ITO-PET was based on the 458 m/z features shared between the two slides, whereas the three-slide comparison was based on the 138 m/z features shared among ITO-glass, ITO-PET, and PEN-glass.

For metabolite annotation, features with signal intensities >100 were retained and annotated using MetaboScape 2025b with a mass tolerance of 15 ppm against the Human Metabolome Database (HMDB) and an in-house reference library provided by Bruker. Metabolite assignments based solely on accurate-mass matching were reported as putative annotations.

### 2.5 Laser Microdissection and Protein Digestion

For proteomic analysis, mouse brain tissue sections mounted on ITO-PET and PEN-glass slides were subjected to laser microdissection (LMD) using an MMI CellCut Plus system (Molecular Machines & Industries, Germany) equipped with a 355 nm solid-state UV laser.^38, 39^ For the comparative evaluation of proteomic performance across the three slides (ITO-glass, ITO-PET, and PEN-glass; Figure 4), regions of interest (ROIs) with an area of 0.2 mm² were collected from each substrate. ITO-PET and PEN-glass slides were microdissected using cutting mode, whereas tissue regions on commercial ITO-glass slides were collected using ablation mode. For evaluation of the effect of MALDI-MSI acquisition on subsequent proteomic analysis using ITO-PET slides (Figure 5), hippocampal regions with sampling areas of 0.05, 0.1, and 0.5 mm² were collected before and after MALDI-MSI, with three replicate measurements performed for each sampling area. For multi-organ validation, a fixed sampling area of 0.1 mm² was used for mouse kidney, lung, spleen, and liver tissues.

For cutting-mode microdissection, the following parameters were used: laser power, 90%; cutting speed, 20 μm/s; objective magnification, 20 ×; laser focus offset, 167 μm; and one cutting repeat. The collected microdissected tissue fragments were subsequently processed for bottom-up proteomic analysis. The detailed procedure was as follows^40, 41^: (1) The microdissected tissue fragments were collected onto the adhesive caps of 0.2 mL MMI Isolation Caps (Molecular Machines & Industries, Germany). (2) A 20 μL aliquot of lysis buffer containing 50 mM NH HCO, 10 mM TCEP, 25 mM IAM, and 0.2% ProteaseMAX Surfactant (Promega, USA) was added to the cap to fully cover the tissue, followed by incubation for 1 hour at room temperature. (3) Proteins were digested by adding a mixture of trypsin (1:10, enzyme-to-protein ratio) and Lys-C (1:10) in 20 μL of 50 mM NH HCO buffer and incubating at 37 °C for 16 hours in a thermomixer. (4) The digest was acidified with formic acid to pH < 4. (5) Peptides were desalted using C18 StageTips, eluted, and concentrated by vacuum centrifugation. (6) The dried peptides were reconstituted in LC-MS grade water containing 0.1% formic acid for subsequent analysis.

### 2.6 LC-MS/MS Analysis

Nanoflow LC-MS/MS analysis was primarily performed using an Orbitrap Exploris 480 mass spectrometer (Thermo Fisher Scientific) coupled to an UltiMate 3000 HPLC system. Peptides were loaded and washed on a self-packed trapping cartridge and subsequently separated on an ACQUITY UPLC HSS T3 analytical column (100 Å, 1.8 μm, 75 μm × 200 mm; Waters) using a 60 min gradient. Solvent B consisted of acetonitrile containing 0.1% formic acid. The gradient was programmed as follows: 3–30% B from 0 to 45 min at 300 nL/min; held at 30% B from 45 to 48 min; increased from 30 to 43% B from 48 to 53 min; increased from 43 to 85% B from 53 to 54 min; held at 85% B from 54 to 58 min at 350 nL/min; decreased from 85 to 3% B from 58 to 59 min at 350 nL/min; and held at 3% B from 59 to 60 min at 300 nL/min, followed by column washing and re-equilibration with solvent A. Peptides were ionized at a spray voltage of 1.8 kV with the capillary temperature maintained at 250 °C. The Orbitrap Exploris 480 was operated in data-independent acquisition (DIA) mode.^42, 43^ Full-scan MS spectra were acquired over an m/z range of 350–1600 at a resolution of 60,000, with a maximum injection time of 50 ms and an automatic gain control (AGC) target of 200%. MS/MS spectra were acquired using higher-energy collisional dissociation (HCD) with a normalized collision energy of 32%. The MS/MS resolution was set to 45,000 over an m/z range of 110–2000, with a maximum injection time of 86 ms and a normalized AGC target of 1000%.

For the supplementary pre- and post-MALDI proteomic analysis of tissue sections mounted on PEN-glass slides, LC-MS/MS analysis was performed using a timsTOF Pro mass spectrometer (Bruker Daltonics, Bremen, Germany) coupled to a nanoElute LC system. Peptides were separated on a 25 cm IonOpticks analytical column maintained at 35 °C at a flow rate of 300 nL/min using a 60 min gradient. The gradient started at 2% solvent B, increased to 22% B at 45 min, 37% B at 50 min, and 80% B at 55 min, and was maintained at 80% B until 60 min. The timsTOF Pro was operated in positive-ion mode with a CaptiveSpray source using dia-PASEF acquisition. The mass range was m/z 100–1700, and the ion mobility range was 0.75–1.30 V s cm ² (1/K), with a TIMS ramp time of 100 ms. The capillary voltage was set to 1500 V, the dry gas flow rate to 3.0 L/min, and the dry temperature to 180 °C. High Sensitivity Detection was disabled. Additional instrument settings included an ion energy of 5 eV, collision energy of 10 eV, transfer time of 60 μs, collision RF of 1500 Vpp, and pre-pulse storage time of 12 μs.

Raw LC-MS/MS data were processed using Spectronaut 19 (Biognosys, Switzerland) with the directDIA workflow. Database searching was performed against the Mus musculus UniProt reference proteome (UP000000589, canonical sequences and isoforms), with Trypsin/P specified as the digestion enzyme. Protein inference and grouping were performed using the algorithms implemented in Spectronaut, and proteins that could not be unambiguously distinguished based on the identified peptide evidence were reported as protein groups. Precursor ion intensities and protein group identifications were used for subsequent quantitative comparisons among the different slides and for evaluating proteomic performance before and after MALDI-MSI.

## 3 Results and Discussion

### 3.1 Characterization of the ITO-PET Slide and Evaluation of MALDI-MSI Performance

Prior to MALDI mass spectrometry imaging, atomic force microscopy (AFM) was employed to characterize the surface morphology and phase homogeneity of the custom-fabricated ITO-PET slide. The three-dimensional topographic image showed a relatively smooth and homogeneous surface morphology (Figure S1A). Quantitative analysis over a 10 × 10 μm² scanning area yielded an average roughness (Ra) of 2.25 nm and a root-mean-square roughness (Rq) of 2.97 nm, with a maximum height variation of 34.0 nm. In addition, AFM phase imaging revealed a highly uniform phase distribution across the scanned region, with phase angles ranging from −71.4° to −73.7°, corresponding to a narrow phase variation of only 2.3° within the 10 × 10 μm² area (Figure S1B). These results indicate that the ITO-PET film possesses a relatively smooth surface and homogeneous interfacial properties, providing a suitable substrate for subsequent MALDI-MSI analysis.

Mouse brain coronal sections were selected as the model tissue for this comparative study due to their well defined anatomical architecture and their widespread use in MALDI mass spectrometry imaging.^44^ The performance of three different substrate materials was systematically evaluated. Analysis of the average mass spectral profiles revealed that the commercial ITO-glass slide and the ITO PET slide produced highly congruent spectral contours and largely overlapping peak patterns (Figure 1A). Both ITO containing substrates exhibited robust lipid signals in the m/z 500–1000 range, indicating comparable desorption/ionization efficiencies. In contrast, the spectral profile of the PEN glass slide was markedly different; the intensities of characteristic lipid peaks were reduced by 60–80% relative to those obtained from the ITO based substrates, suggesting substantially compromised ionization efficiency. As shown in Figure 1B, Venn diagram visualization further quantified the similarity in detected m/z features among the three slides. For calculation of the percentages shown in the Venn diagram, the union of all unique m/z features detected across ITO-glass, ITO-PET, and PEN-glass was defined as 100%, and the percentage of each region was calculated relative to this total union. The m/z features shared between ITO-glass and ITO-PET accounted for 56.7% of the total union, reflecting substantial consistency between the two conductive substrates. In contrast, the shared features involving PEN-glass accounted for less than 10% of the total union, indicating substantially lower concordance with either ITO-based substrate. This pronounced difference may be attributable, at least in part, to the insufficient surface conductivity of PEN-glass, which can adversely affect MALDI ion generation.^45, 46^

**Figure 1.**
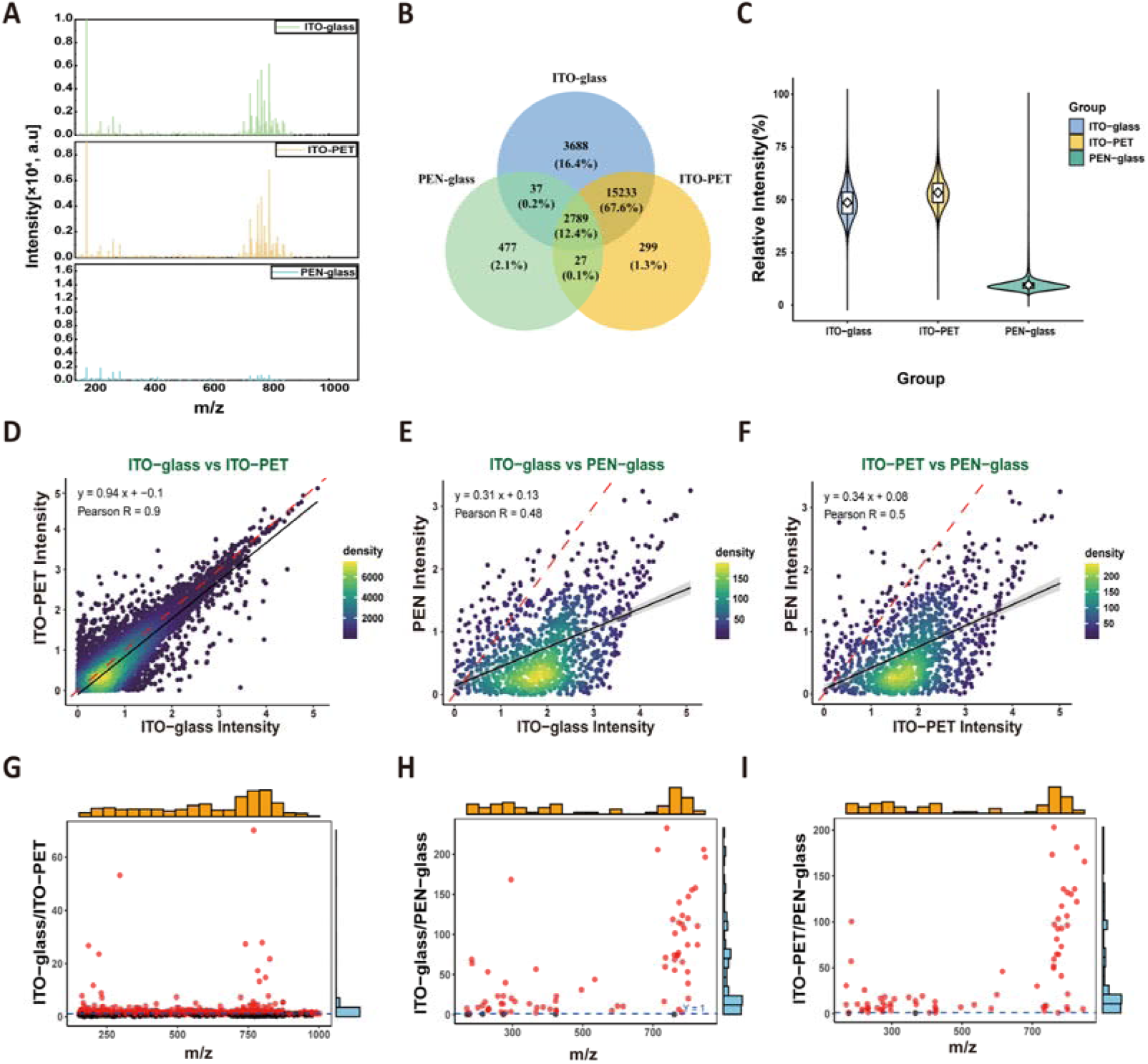
Comparative evaluation of MALDI-MS signal characteristics among ITO-glass, ITO-PET, and PEN-glass slides. (A) Average mass spectra obtained from mouse brain tissue sections on the three slides. (B) Venn diagram showing the overlap of detected m/z features among ITO-glass, ITO-PET, and PEN-glass. Numbers indicate the counts of unique or shared m/z features in each region, with the corresponding percentages shown in parentheses. (C) Distribution of normalized MALDI-MS signal intensities obtained from mouse brain tissue sections on the three slides. Violin plots show the distributions of relative signal intensities (%), with embedded boxplots indicating the median and interquartile range (IQR); white diamonds represent the mean values. The mean relative signal intensities were 48.65% for ITO-glass, 53.36% for ITO-PET, and 9.64% for PEN-glass. (D–F) Pairwise correlation analysis of signal intensities for shared m/z features detected on different slides: (D) ITO-glass versus ITO-PET (Pearson’s R = 0.90), (E) ITO-glass versus PEN-glass (R = 0.48), and (F) ITO-PET versus PEN-glass (R = 0.50). (G–I) Group marginal plots of pairwise intensity ratios for shared m/z features with signal intensities >50, illustrating the m/z regions in which ions with enhanced signal intensities were predominantly distributed: (G) ITO-glass/ITO-PET, (H) ITO-glass/PEN-glass, and (I) ITO-PET/PEN-glass.

The distributions of normalized ion signal intensities were further compared among the three slides (Figure 1C). For each slide, ion intensities were normalized to the maximum detected intensity, which was defined as 100%, and expressed as relative intensity (%). The resulting intensity distributions were visualized using violin plots. ITO-glass and ITO-PET exhibited broadly similar intensity distributions, whereas PEN-glass displayed a markedly lower intensity profile. The mean relative signal intensities were 48.65% for ITO-glass, 53.36% for ITO-PET, and 9.64% for PEN-glass. The difference between ITO-glass and ITO-PET was only 4.71 percentage points, markedly smaller than the differences between ITO-glass and PEN-glass (39.02 percentage points) and between ITO-PET and PEN-glass (43.72 percentage points). Together, these data indicate that the signal intensity characteristics of ITO-PET closely resembled those of conventional ITO-glass, while clearly differing from the substantially lower intensities observed with PEN-glass.

Using MALDI MSI data, the abundance of ions at identical m/z values was quantitatively evaluated across three types of slides: commercial ITO glass, the ITO PET slide, and commercial PEN glass. All detected MS signals with intensities >1 were log-transformed prior to Pearson correlation analysis^47^ and linear regression modeling^47, 48^ for pairwise comparison (Figures 1D-F). Statistical analysis revealed excellent concordance between the ITO glass and the ITO-PET slides in detecting ions at identical m/z values, with a Pearson correlation coefficient of R = 0.9. The concentration of data points around the regression line indicates a high degree of consistency in the relative ion signal patterns obtained from the two slides. In contrast, comparative analyses between each ITO-based substrate and the PEN-glass (Figures 1E and 1F) revealed substantially fewer matched m/z ions, lower correlation coefficients, and a markedly dispersed distribution of data points. Building upon these observations, a more detailed analysis of ion signal intensity ratios was performed in figure 1G-I. Mass spectrometry data were processed by retaining only ions with an intensity greater than 50. For shared peaks, intensity ratios between the custom-fabricated ITO-PET slide and the commercial ITO glass slide were calculated. The comparison revealed no consistent intensity advantage for either substrate, with some ions showing higher intensities on the ITO-PET slide and others on the ITO glass. These ions spanned a broad range of m/z values but were predominantly distributed within the lipid region. Notably, when ITO-glass and ITO-PET were compared with PEN-glass, the majority of shared m/z features exhibited higher signal intensities on the two ITO-based slides than on PEN-glass. Moreover, the response profiles of these ions varied across the m/z range, with signal gaps observed in specific intervals in Figures 1H and 1I. Collectively, these findings demonstrate the substantially reduced performance of non-conductive PEN-glass in MALDI-MSI and highlight the importance of adequate surface conductivity for reliable and reproducible spatial metabolomic measurements.

### 3.2 Comparison of Slide Performance via Spatial Clustering and Dimensionality Reduction

To evaluate the spatial organization of MALDI-MSI data acquired from different slides, Leiden clustering was applied to the pixel-wise mass spectra of mouse brain sections using consistent clustering parameters across all datasets (Figures 2A–C). The commercial ITO-glass and custom-fabricated ITO-PET slides yielded 27 and 26 spatial clusters, respectively, both exhibiting coherent spatial organization and relatively well-defined regional boundaries. Notably, the hippocampal region showed clear and continuous spatial partitioning on both substrates, with its overall morphology and regional integrity well preserved. The clustering pattern obtained with ITO-PET closely resembled that observed with conventional ITO-glass, particularly in the hippocampal region, although some local boundaries appeared slightly less distinct. PEN-glass yielded 38 clusters; however, rather than the number of clusters itself, the spatial organization of these clusters differed noticeably from that observed with the two ITO-based substrates. Specifically, the PEN-glass-derived clusters exhibited more fragmented and dispersed spatial distributions, particularly in the hippocampal region, where regional boundaries were less distinct and spatial continuity was reduced. Together with the substantially lower signal intensities observed for PEN-glass, these results suggest that the reduced signal quality obtained with this substrate may contribute to decreased spatial coherence in the resulting molecular clustering.

**Figure 2.**
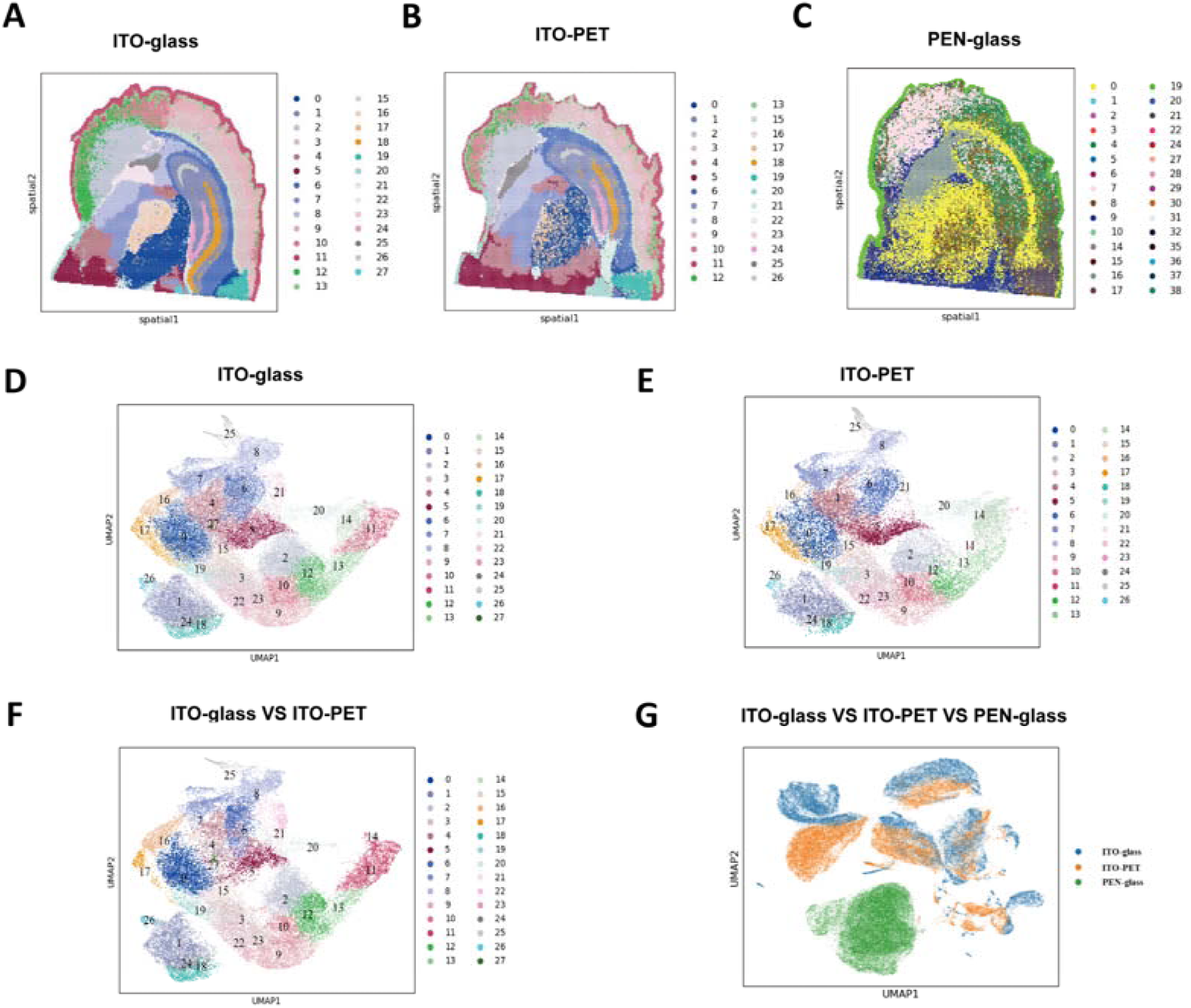
Comparison of spatial clustering and high-dimensional spectral characteristics of MALDI-MSI data acquired from ITO-glass, ITO-PET, and PEN-glass slides. (A–C) Leiden clustering maps of pixel-wise MALDI-MSI spectra obtained from mouse brain tissue sections on (A) ITO-glass, (B) ITO-PET, and (C) PEN-glass slides, yielding 27, 26, and 38 spatial clusters, respectively. (D–E) UMAP projections of pixel-wise spectral data obtained from (D) ITO-glass and (E) ITO-PET slides, with pixels colored according to the corresponding Leiden-derived clusters. (F) Combined UMAP projection of pixels from ITO-glass and ITO-PET slides based on the 458 m/z features shared between the two slides. (G) Combined UMAP projection of pixels from ITO-glass, ITO-PET, and PEN-glass slides based on the 138 m/z features shared among all three slides.

To further compare the high-dimensional spectral characteristics of the three slides, uniform manifold approximation and projection (UMAP) was applied to the pixel-wise spectral data (Figures 2D–G). Individual UMAP projections of ITO-glass and ITO-PET showed similarly organized spectral populations corresponding to the Leiden-derived clusters (Figures 2D and 2E). When the analysis was restricted to the 458 m/z features shared between ITO-glass and ITO-PET, pixels from the two slides showed substantial overlap in UMAP space (Figure 2F), indicating a high degree of similarity in their multivariate spectral characteristics. When UMAP analysis was performed using the 138 m/z features shared among all three slides, PEN-glass pixels occupied a more distinct region of the UMAP space, whereas the ITO-glass and ITO-PET populations showed greater overlap (Figure 2G). These observations indicate that, under the experimental and data-processing conditions used here, the spectral characteristics obtained with ITO-PET more closely resembled those obtained with conventional ITO-glass than those obtained with PEN-glass.

Collectively, the Leiden clustering and UMAP analyses demonstrate that ITO-PET supports coherent spatial molecular patterns and multivariate spectral characteristics closely comparable to those obtained with conventional ITO-glass. In particular, the spatial continuity and regional organization of the hippocampus were well preserved on both ITO-based substrates. In comparison, PEN-glass exhibited more fragmented spatial clustering and a more distinct distribution in the reduced-dimensional spectral space, consistent with its lower overall MALDI-MS signal performance.

### 3.3 Comparison of Slide Performance via Metabolite Annotation and Spatial Molecular Distributions

To evaluate the performance of the three slides—commercial ITO-glass, custom-fabricated ITO-PET, and commercial PEN-glass—for spatial metabolomic analysis, we compared their metabolite annotation coverage and the spatial distributions of characteristic molecules. All MALDI-MSI datasets were processed using a uniform peak-intensity threshold of >100 during peak extraction to ensure consistent data processing across the three slides. Under these conditions, 357, 386, and 118 putatively annotated metabolites were obtained from ITO-glass, ITO-PET, and PEN-glass, respectively (Figure 3A), with 46 metabolites commonly annotated across all three slides (Figure 3B). The comparable numbers of annotated metabolites obtained from ITO-glass and ITO-PET, together with the substantially lower number detected on PEN-glass, indicate similar metabolite detection coverage between the two ITO-based slides.

**Figure 3.**
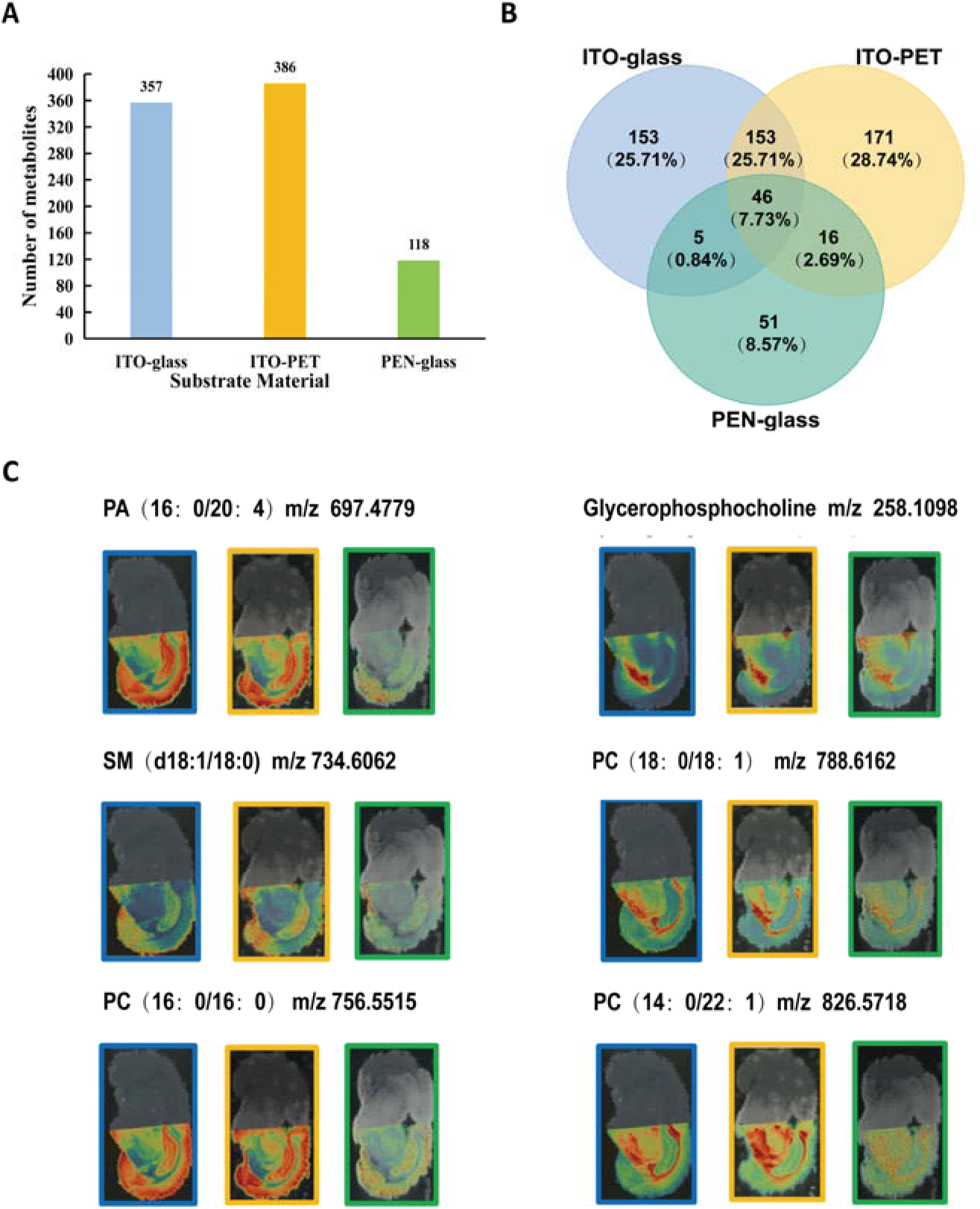
Comparative evaluation of metabolite annotation coverage and spatial molecular distributions across ITO-glass, ITO-PET, and PEN-glass slides. (A) Numbers of putatively annotated metabolites obtained from mouse brain tissue sections on the three slides using a uniform peak-intensity threshold of >100 during feature extraction. A total of 357, 386, and 118 metabolites were annotated from ITO-glass, ITO-PET, and PEN-glass, respectively. (B) Venn diagram showing the overlap of putatively annotated metabolites among the three slides, with 46 metabolites commonly annotated across all three slides. (C) Representative MALDI-MSI ion images showing the spatial distributions of six characteristic metabolites in mouse brain tissue sections across the three slides. PA(16:0/20:4), SM(d18:1/18:0), and PC(16:0/16:0) exhibited relatively high signal intensities in the hippocampal region, whereas glycerophosphocholine, PC(18:0/18:1), and PC(14:0/22:1) exhibited relatively low signal intensities in this region. ITO-glass and ITO-PET showed similar spatial distribution patterns with clear regional contrast, whereas the corresponding ion images obtained on PEN-glass exhibited weaker signals and less distinct regional distributions.

To further compare the spatial distributions of characteristic molecules, six representative metabolites exhibiting distinct distribution patterns in the hippocampal region were examined (Figure 3C). PA(16:0/20:4), SM(d18:1/18:0), and PC(16:0/16:0) showed pronounced enrichment in the hippocampal region, whereas glycerophosphocholine, PC(18:0/18:1), and PC(14:0/22:1) exhibited relatively low abundance in this region. The ion images acquired using ITO-glass and ITO-PET showed highly similar spatial distribution patterns for these characteristic molecules, with clear regional contrast and well-defined localization. In particular, the characteristic enrichment and low-abundance patterns in the hippocampal region were consistently visualized on both ITO-based slides. In contrast, the corresponding ion images obtained using PEN-glass exhibited substantially weaker signals and less distinct regional distribution patterns.

Collectively, these results demonstrate that the ITO-PET slide provides metabolite detection coverage and spatial distributions of characteristic molecules closely comparable to those obtained with the conventional ITO-glass slide. In contrast, the PEN-glass slide yielded substantially fewer annotated metabolites and markedly weaker ion signals, with the characteristic molecular distribution patterns being less clearly visualized. These findings further support the suitability of the ITO-PET slide for MALDI-MSI-based spatial metabolomic analysis.

### 3.4 Comparative Evaluation of Proteomic Performance across Different Slides

The performance of the three slides for spatially resolved proteomics was evaluated following laser microdissection (Figure S2A–C) and subsequent sample processing. As shown in Figure 4A, the distributions of log10-transformed proteomic signal intensities were compared after excluding zero-intensity values. The ITO-PET and PEN-glass slides exhibited highly comparable intensity distributions, with mean log10 intensities of 4.00 and 4.03, respectively, whereas the ITO-glass slide showed a lower overall intensity distribution, with a mean value of 3.24. These results indicate that the overall proteomic signal intensity obtained with ITO-PET was closely comparable to that obtained with the conventional PEN-glass slide and higher than that observed with ITO-glass. To further evaluate the consistency of the proteomic profiles, pairwise correlation analyses were performed based on the intensities of protein groups commonly identified between the compared slides. A strong correlation was observed between ITO-PET and PEN-glass (Pearson’s R = 0.94; Figure 4B), whereas ITO-glass and PEN-glass also showed a high, although slightly lower, correlation (R = 0.91; Figure 4C). These results demonstrate a high degree of consistency in protein group abundance profiles among the slides, with ITO-PET showing particularly close agreement with the conventional PEN-glass slide.

**Figure 4.**
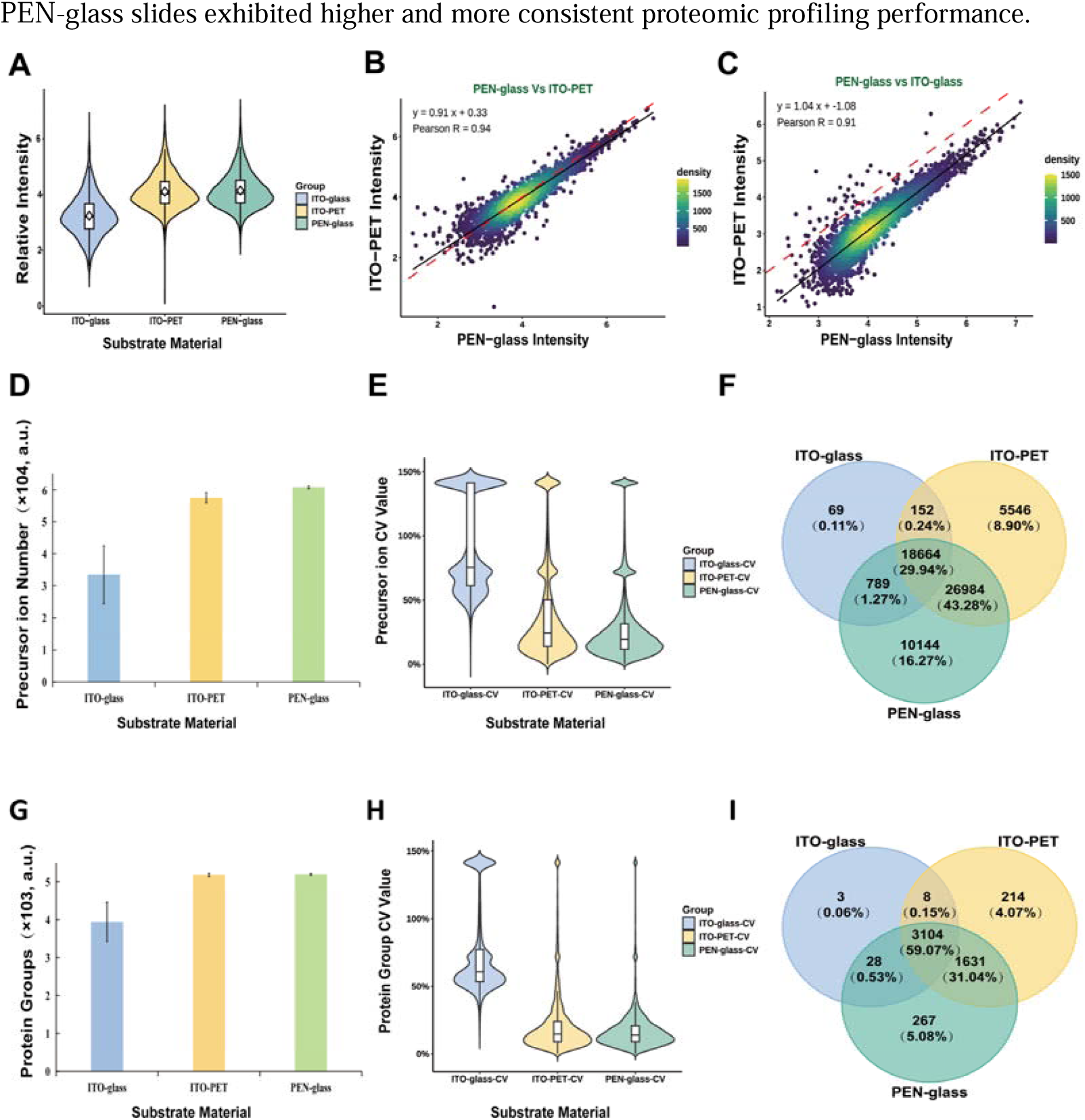
Comparative evaluation of proteomic performance across three slides. (A) Distribution of log10-transformed protein group intensities obtained from mouse brain tissue sections collected from ITO-glass, ITO-PET, and PEN-glass slides. Zero-intensity values were excluded before analysis. Violin plots show the distributions of log10-transformed protein group intensities, with embedded boxplots indicating the median and interquartile range (IQR); white diamonds represent the mean values. The mean log10 protein group intensities were 3.24 for ITO-glass, 4.00 for ITO-PET, and 4.03 for PEN-glass. (B–C) Scatter plots showing pairwise correlations of the intensities of protein groups commonly identified between the compared slides: (B) PEN-glass versus ITO-PET and (C) PEN-glass versus ITO-glass. (D) Bar plot showing the numbers of precursor ions detected from the three slides. (E) Violin plot showing the distributions of coefficients of variation (CVs) for precursor ion intensities across replicate measurements for the three slides. (F) Venn diagram showing the overlap of precursor ions detected across the three slides. (G) Bar plot showing the numbers of protein groups identified from the three slides. (H) Violin plot showing the distributions of coefficients of variation (CVs) for protein group intensities across replicate measurements for the three slides. (I) Venn diagram showing the overlap of protein groups identified across the three slides.

Quantitative comparison of precursor ion and protein group identifications further showed higher proteomic yields for ITO-PET and PEN-glass than for ITO-glass under the sampling conditions used. The ITO-PET and PEN-glass slides yielded highly comparable numbers of precursor ions and identified protein groups (Figure 4D and 4G), indicating consistent and robust proteomic profiling performance. In contrast, the ITO-glass slide showed lower numbers of both precursor ions and identified protein groups than the ITO-PET and PEN-glass slides, indicating a reduced capacity for proteomic profiling under the laser ablation mode used for tissue collection. These results indicate more favorable downstream proteomic performance for ITO-PET and PEN-glass under the conditions evaluated.

To further assess quantitative reproducibility, the coefficients of variation (CVs) of precursor ion and protein groups intensities were calculated across replicate measurements. As shown in Figures 4E and 4H, the ITO-glass slide exhibited a tendency toward higher CV values for both precursor ion and protein group intensities, indicating greater variability in the measurements. In comparison, ITO-PET and PEN-glass, both of which are compatible with cutting-mode microdissection, showed generally lower CV distributions for precursor ion and protein group intensities. These descriptive results suggest that cutting-mode tissue collection using ITO-PET and PEN-glass may provide more consistent quantitative proteomic measurements than ablation-mode collection from ITO-glass.

Venn diagram visualization was subsequently performed to compare the overlap of precursor ion and protein groups identified across the three slides (Figure 4F and 4I). The ITO-PET and PEN-glass slides showed substantial overlap in both precursor ions and identified protein groups, with overlap rates of 73.46% and 90.26%, respectively, indicating a high degree of similarity in their proteomic feature profiles. In contrast, the overlap between the ITO-glass slide and either the ITO-PET or PEN-glass slide was below 60% for both precursor ions and identified protein groups. Notably, despite these relatively lower overlap rates, the shared precursor ions and protein groups accounted for the majority of the features identified on the ITO-glass slide. Together with the substantially fewer precursor ions and protein group identified on the ITO-glass slide, these results indicate a more limited proteomic profiling capacity under ablation-mode sampling, whereas the ITO-PET and PEN-glass slides exhibited higher and more consistent proteomic profiling performance.

### 3.5 Validation of MALDI-MSI Workflow Compatibility with Downstream Proteomic Analysis

To determine whether MALDI-MSI acquisition affects subsequent proteomic analysis of the same tissue section, we first evaluated proteomic performance before and after MALDI-MSI using conventional PEN-glass slides commonly employed for laser microdissection. As shown in Figure S3A–D, the overall proteomic signal intensity distributions, precursor ion counts, and protein group identifications were broadly comparable before and after MALDI-MSI. These descriptive comparisons suggest that MALDI-MSI acquisition had limited effects on subsequent proteomic measurements under the experimental conditions evaluated.

We next evaluated the compatibility of the ITO-PET slide with downstream proteomic analysis following MALDI-MSI acquisition. Hippocampal regions of mouse brain tissue were collected at three sampling areas (0.05, 0.1, and 0.5 mm²), and proteomic analyses were performed before and after MALDI-MSI (Figure 5A). Overall proteomic signal intensities showed similar distributions between the pre- and post-MALDI groups at each sampling area (Figure 5B). Precursor ion counts and protein group identifications showed modest decreases after MALDI-MSI at the three sampling areas, while their overall levels remained comparable between the pre- and post-MALDI groups (Figures 5C and 5D).

**Figure 5.**
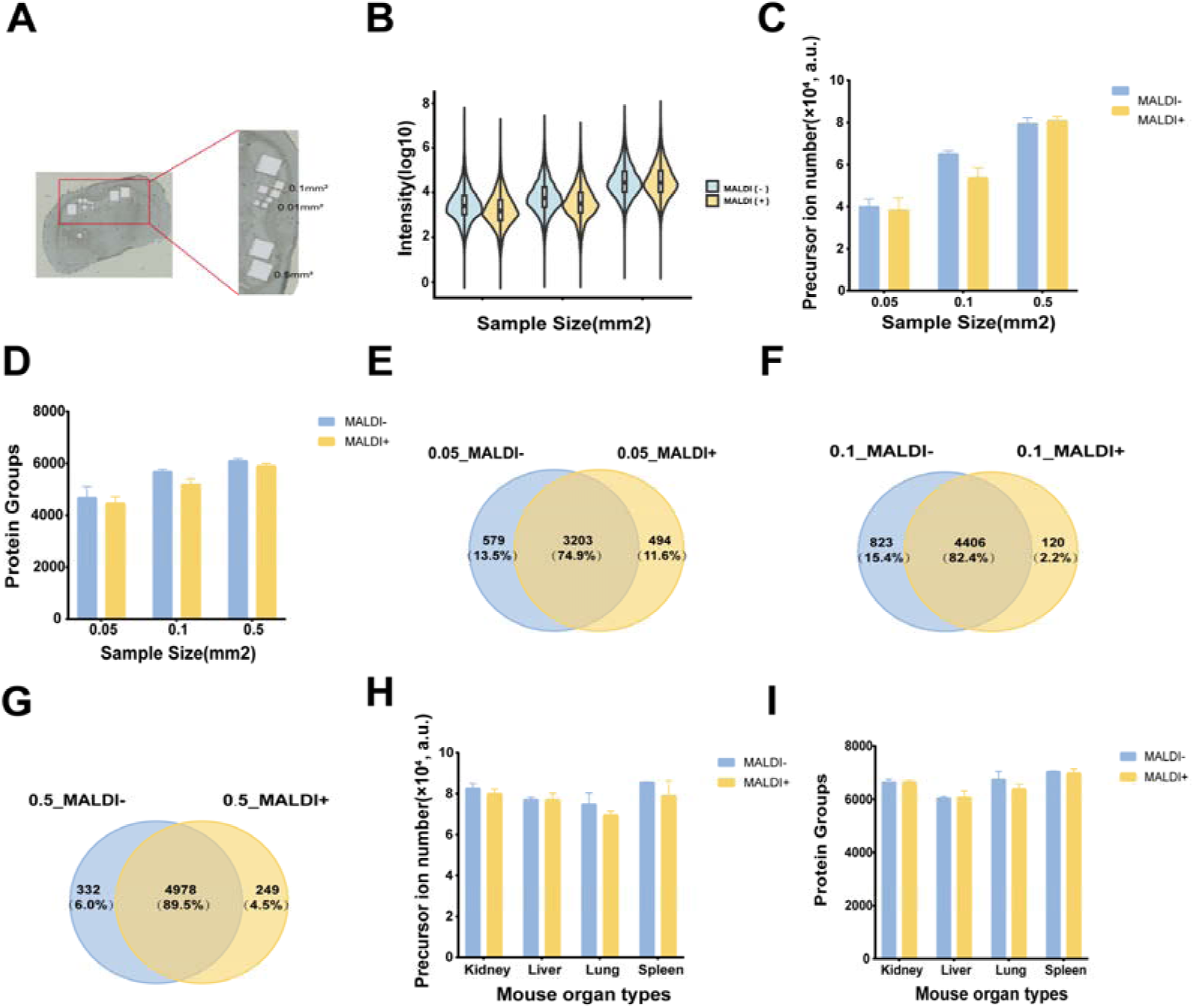
Evaluation of the effect of MALDI-MSI on downstream proteomic analysis across different tissue sampling areas and organs. (A) Representative scanning images of mouse brain sections after microdissection, showing hippocampal regions collected at three microdissection areas (0.05, 0.1, and 0.5 mm²), with three replicates analyzed for each area. (B) Violin plots comparing overall proteomic signal intensities (log10-transformed) before and after MALDI-MSI across the three microdissection areas. (C) Bar charts comparing precursor ion counts before and after MALDI-MSI across the three microdissection areas. (D) Bar charts comparing protein group identification numbers before and after MALDI-MSI across the three microdissection areas. (E) Venn diagram showing the overlap of identified protein groups before and after MALDI-MSI for a microdissection area of 0.05 mm². (F) Venn diagram showing the overlap of identified protein groups before and after MALDI-MSI for a microdissection area of 0.1 mm². (G) Venn diagram showing the overlap of identified protein groups before and after MALDI-MSI for a microdissection area of 0.5 mm². (H) Bar charts comparing precursor ion counts before and after MALDI-MSI across mouse kidney, liver, lung, and spleen tissues collected using a microdissection area of 0.1 mm². (I) Bar charts comparing protein group identification numbers before and after MALDI-MSI across mouse kidney, liver, lung, and spleen tissues collected using a microdissection area of 0.1 mm².

Venn diagram visualization further showed substantial overlap in protein groups identified before and after MALDI-MSI (Figures 5E–G). For each comparison, the union of all protein groups identified in the pre- and post-MALDI groups was defined as 100%, and the percentage of shared protein groups was calculated relative to this union. The shared protein groups accounted for 74.9%, 82.4%, and 89.5% of the total union for sampling areas of 0.05, 0.1, and 0.5 mm², respectively. The proportion of shared protein groups increased with increasing sampling area, with the highest concordance observed at 0.5 mm². Together with the similar signal intensity distributions and comparable identification numbers, these descriptive results indicate that the overall proteomic profiles were largely preserved following MALDI-MSI acquisition across the tissue sampling areas examined.

To further assess the applicability of the workflow across different tissue types, parallel experiments were performed using mouse kidney, lung, spleen, and liver tissues with a fixed microdissection area of 0.1 mm² (Figure S4A–D). Venn diagram visualization showed substantial overlap in protein groups identified before and after MALDI-MSI, with shared protein groups accounting for 95.3% of the total union for kidney, 89.5% for lung, 93.3% for spleen, and 90.6% for liver (Figure S5A–D). Precursor ion counts (Figure 5H) and protein group identifications (Figure 5I) were also broadly comparable before and after MALDI-MSI across the four organs examined. These multi-organ observations were consistent with those obtained from mouse brain tissue and further support the compatibility of MALDI-MSI acquisition with subsequent proteomic analysis across the tissue types evaluated.

Collectively, these descriptive comparisons indicate that MALDI-MSI acquisition had limited effects on subsequent proteomic measurements using both conventional PEN-glass and custom-fabricated ITO-PET slides under the experimental conditions evaluated. The substantial overlap in protein group identifications, together with the broadly comparable proteomic signal profiles and identification numbers before and after MALDI-MSI, supports the feasibility of sequential spatial metabolomic and proteomic analysis from the same tissue section using the ITO-PET slide.

## 4 Conclusions

In this study, we developed and systematically evaluated a conductive ITO-PET slide for sequential MALDI-MSI-based spatial metabolomics and LCM-based proteomics from the same tissue section. The slide was designed to address the different material requirements of the two analytical modalities by combining the electrical conductivity required for MALDI-MSI with the mechanical properties needed for cutting-mode laser microdissection. This design provides an alternative to conventional ITO-glass, which requires ablation-mode tissue collection for downstream proteomics, and PEN-glass, which is compatible with LCM cutting but provides substantially lower MALDI-MS signal performance under the conditions evaluated.

Using mouse brain tissue, the ITO-PET slide showed MALDI-MSI performance closely comparable to conventional ITO-glass in terms of spectral characteristics, ion detection coverage, signal intensity distribution, metabolite annotation, and preservation of spatial molecular patterns. For downstream proteomic analysis, cutting-mode tissue collection from ITO-PET yielded proteomic signal intensities and identification numbers comparable to those obtained with conventional PEN-glass, together with generally lower CV distributions than those observed for ablation-mode collection from ITO-glass. Evaluation of proteomic measurements before and after MALDI-MSI across different tissue sampling areas showed broadly comparable signal intensity distributions, precursor ion counts, and protein group identifications, together with substantial overlap in identified protein groups. Similar patterns were observed in mouse kidney, lung, spleen, and liver tissues, supporting the applicability of the workflow across the tissue types examined.

More broadly, the ability to obtain complementary molecular information from the same tissue section provides an opportunity to maximize information recovery from spatially limited biological specimens while maintaining close spatial correspondence across molecular layers. This capability may be particularly valuable for highly heterogeneous tissues, where molecular differences between adjacent sections can complicate cross-modal comparisons. With further improvements in analytical sensitivity and spatial resolution, the workflow may enable more comprehensive characterization of spatially coordinated metabolic and proteomic processes and contribute to a deeper understanding of molecular organization and functional heterogeneity within complex tissues.

## Supporting Information

Additional AFM characterization of the ITO-PET film, laser microdissection images, evaluation of proteomic performance before and after MALDI-MSI on PEN-glass slides, multi-organ MALDI-MSI images, and protein group identification overlap analyses (PDF).

## CRediT authorship contribution statement

Haizhu Wang, Yan Li, and Peng Xue conceived the study and designed the experimental strategy and analytical workflow. Haizhu Wang performed the major experiments, including MALDI mass spectrometry imaging, laser microdissection, and LC-MS-based proteomic analysis, and conducted data processing and analysis. Shengmei Liu, Lixian Lin, Yongsheng Cheng, and Yuchuang Zheng contributed to experimental work, data analysis, and interpretation of the results. Yan Li and Peng Xue supervised the study and provided guidance on experimental design, methodology, and data interpretation. Haizhu Wang wrote the original draft. All authors reviewed and revised the manuscript, discussed the results, and approved the final version.

## Data availability

Data will be made available on request.

## Acknowledgments

We gratefully acknowledge Pu Zhang from Guangzhou Laboratory for valuable guidance and assistance with the animal experiments. We also thank Linyuan Fan from Guangzhou Laboratory for helpful guidance and suggestions on manuscript preparation.

