## Additional AFM characterization of the ITO-PET film, laser microdissection images, evaluation of proteomic performance before and after MALDI-MSI on P for "Integrated Spatial Metabolomics and Proteomics from the Same Tissue Section Using a Conductive ITO-PET Slide" (from another version of this preprint: version 1, 2026-09-14)

**Supplementary materials**

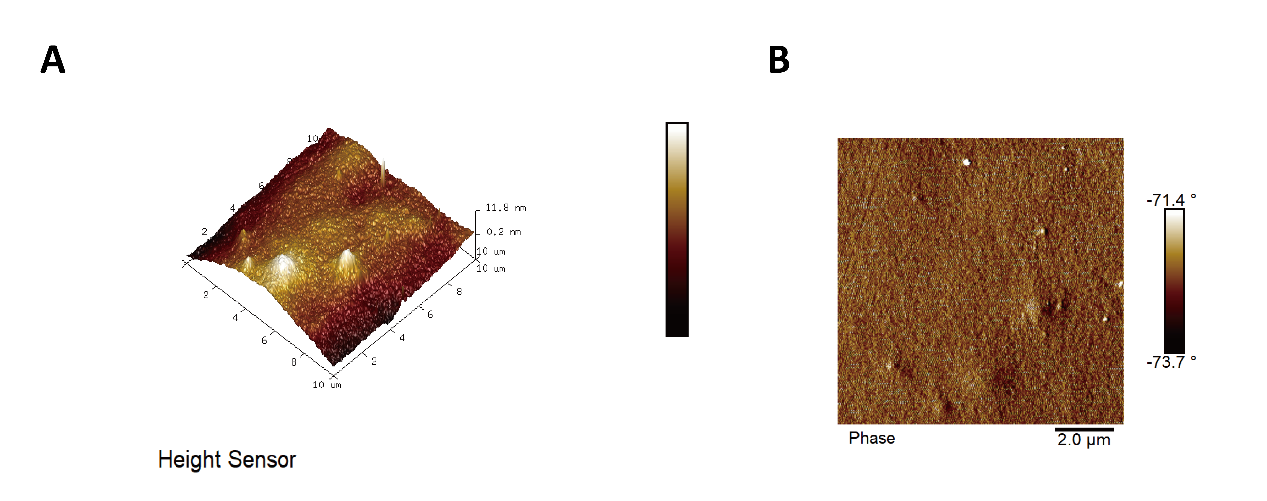

Figure S1. AFM characterization of the surface morphology and phase homogeneity of the ITO-PET film. (A) Three-dimensional AFM topographic image of the ITO-PET film acquired over a 10 × 10 μm² scanning area. The surface exhibited a relatively smooth morphology, with an average roughness (Ra) of 2.25 nm and a root-mean-square roughness (Rq) of 2.97 nm. (B) AFM phase image acquired over a 10 × 10 μm² scanning area, showing a relatively uniform phase distribution. The phase angle ranged from −71.4° to −73.7°, corresponding to a total phase variation of approximately 2.3°.

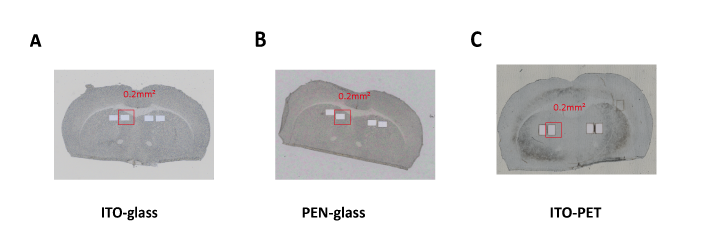

Figure S2. Representative images of laser-microdissected mouse brain tissue sections mounted on ITO-glass, PEN-glass, and ITO-PET slides. (A) Scanning image of microdissected frozen mouse sections mounted on the ITO-glass slide; (B) Scanning image of microdissected frozen mouse sections mounted on the PEN-glass membrane slide; (C) Scanning image of microdissected frozen mouse sections mounted on the ITO-PET membrane slide.

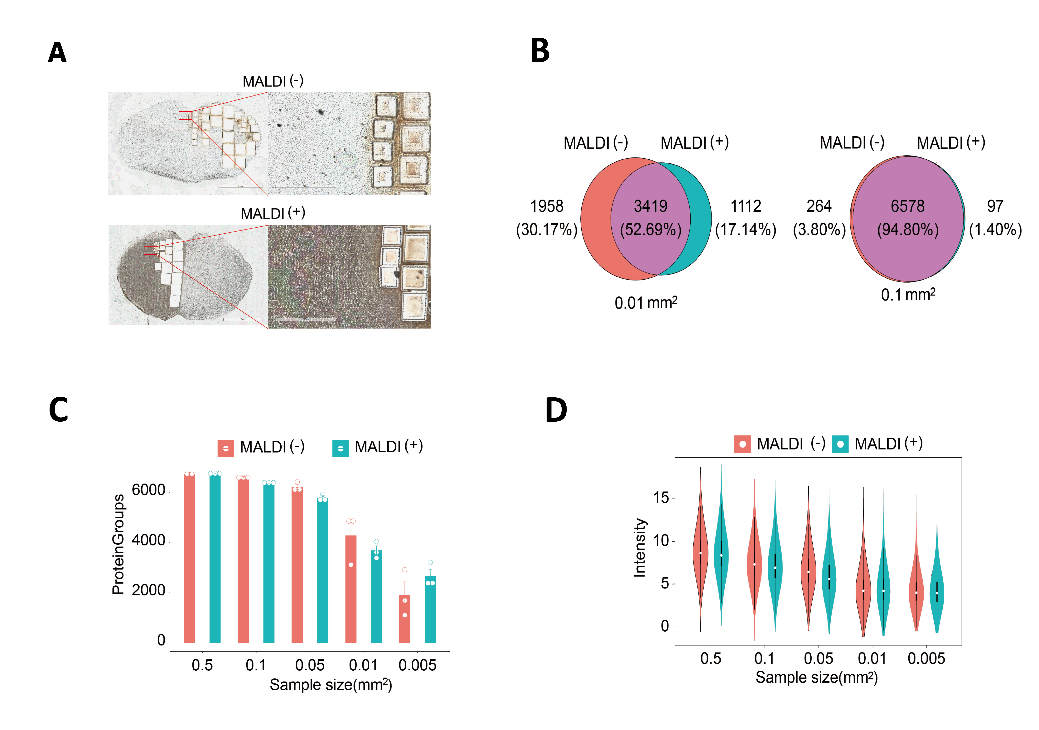

Figure S3. Evaluation of proteomic performance before and after MALDI-MSI on PEN-glass slides across different tissue sampling areas. (A) Representative microscopic images of frozen mouse brain coronal sections mounted on PEN-glass slides before (top) and after (bottom) MALDI-MSI. (B) Venn diagrams showing the overlap of identified protein groups before and after MALDI-MSI at tissue sampling areas of 0.01 and 0.1 mm². The overlap rates were 52.69% and 94.80% for 0.01 and 0.1 mm², respectively. (C) Bar plots comparing the numbers of identified protein groups before and after MALDI-MSI at tissue sampling areas of 0.005, 0.01, 0.05, 0.1, and 0.5 mm². (D) Violin plots comparing the distributions of protein signal intensities before and after MALDI-MSI across the same five tissue sampling areas.

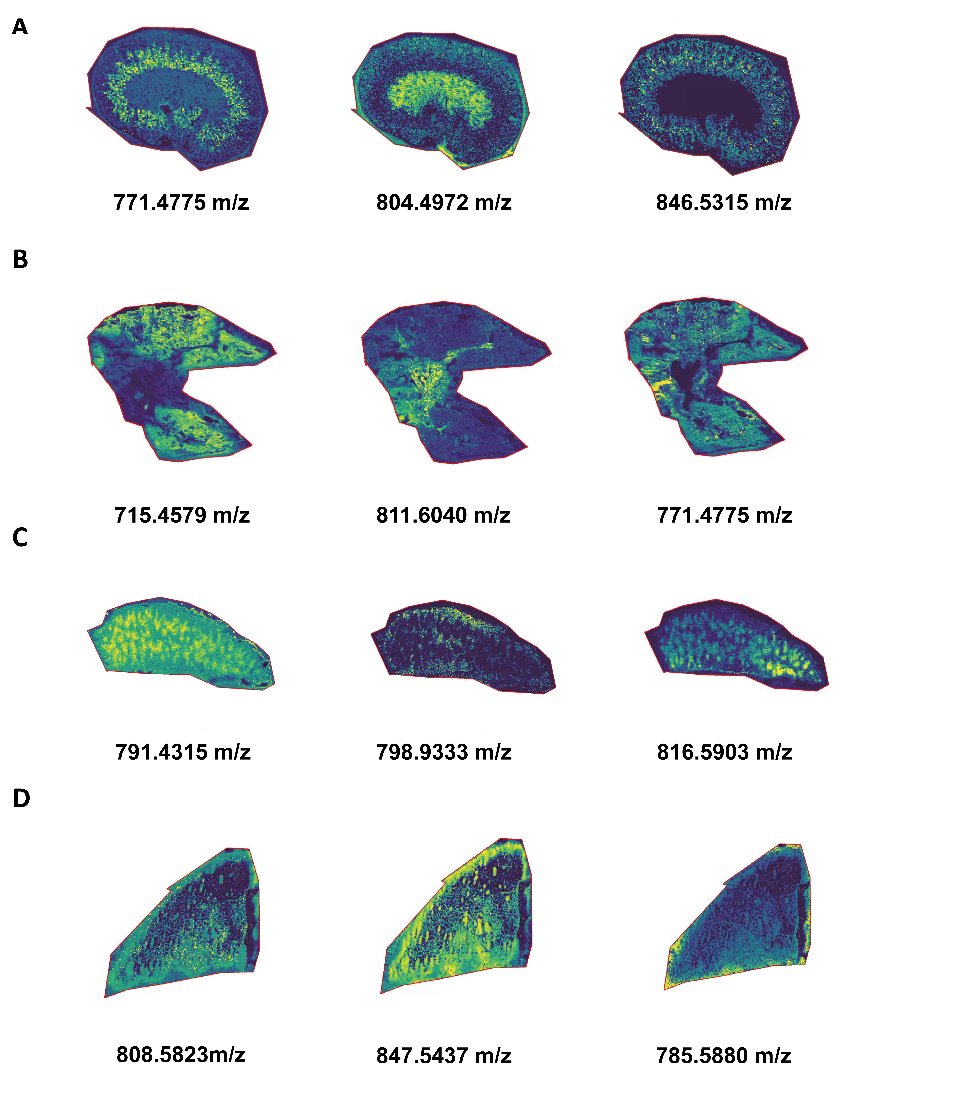

Figure S4. Representative MALDI mass spectrometry imaging (MSI) of metabolites at different mass-to-charge ratios (m/z) in mouse tissues. (A) Kidney; (B) Lung; (C) Spleen; (D) Liver. For each organ, multiple m/z values are depicted to illustrate the spatial heterogeneity and organ-specific metabolic profiles. The intensity scale (from low to high) is shown by the color gradient.

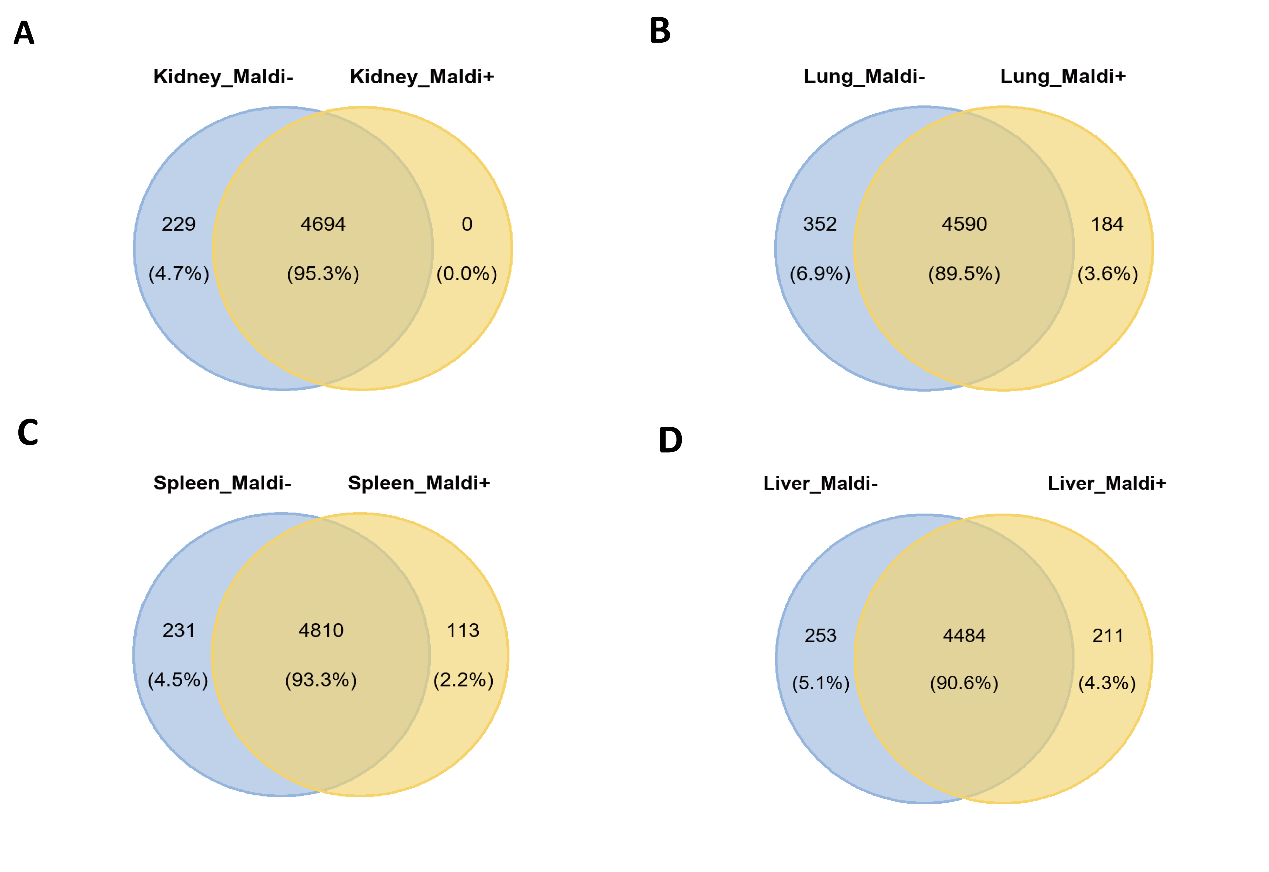

Figure S5. Overlap of protein groups identified before and after MALDI-MSI across different mouse organs. (A–D) Venn diagrams comparing protein groups identified before and after MALDI-MSI from the same tissue sections using a fixed microdissection area of 0.1 mm²: (A) kidney, (B) lung, (C) spleen, and (D) liver. The corresponding overlap percentages are indicated in the Venn diagrams.
